# Accelerated APOA-II senile amyloidosis in a *PAI-1* (plasminogen activator inhibitor 1) knock-out model

**DOI:** 10.64898/2025.12.18.695159

**Authors:** G. Roussine Codo, Pauline Duchatelet, Gemma Rivas-Martinez, Alessio Lampis, Karolina Swiderska, Emilie Pinault, Camille Cohen, Magali Colombat, Sebastien Bender, Sirac Christophe

## Abstract

Apolipoprotein A-II amyloidosis (AApoA-II) is the most common form of age-related amyloidosis in mice, typically arising sporadically in aged animals or in senescence-accelerated strains carrying the APOA2C variant. Here, we report that plasminogen activator inhibitor-1 deficient (PAI-1^−/−^) mice develop systemic AApoA-II amyloidosis with complete penetrance at an intermediate age. Congo red-positive deposits were consistently observed after 12 months in multiple organs, including kidney, spleen, liver, heart, tongue, and intestine, whereas wild-type controls displayed significantly lower amyloid burden and penetrance. Injection of amyloid enhancing factor further accelerated disease onset. Mass spectrometry of purified fibrils and immunofluorescence identified APOA2 as the causative protein. Functional assessment revealed a strong correlation between renal amyloid burden and albuminuria, establishing urinary albumin as a reliable biomarker of disease evolution for preclinical studies. This model provides a robust and reproducible system for investigating age-related amyloidosis and for pre-clinical evaluation of anti-amyloid therapies. Although the role of PAI-1 deficiency in other forms of amyloidosis remains to be demonstrated, our findings raise concern that chronic PAI-1 inhibition, currently pursued in clinical programs, may inadvertently facilitate amyloid deposition by altering extracellular matrix remodeling. PAI-1^−/−^ mice thus represent both a valuable tool for amyloidosis research and a cautionary signal regarding therapeutic strategies targeting PAI-1.

## Introduction

Unlike in humans, where apolipoprotein A-II amyloidosis (AApoAII) is a rare hereditary form, AApoAII is common in mice and considered as the most frequent amyloidosis type in aged animals. Especially, an amyloidogenic variant of APOA2 (*APOA2*^*C*^) has been identified in senescence accelerated mice (SAM), which develop severe systemic amyloidosis from 6 months of age [1]. Beyond this specific variant of APOA2, senile amyloidosis can also occur sporadically in certain mouse genetic backgrounds in old animals. In all cases, the use of these models may be biased by comorbidities related either to age or to the accelerated senescence phenotype observed in SAM mice. Hereby, we show that plasminogen-activator inhibitor knock-out mice (PAI-1^−/−^) present with an accelerated age-related APOA2 amyloidosis.

Originally developed by Carmeliet et al. [2], this model features a deletion of *SERPINE1*, the PAI-1-encoding gene, and is characterized by a mild hyperfibrinolytic state without compromising overall hemostasis. Beyond its role in hemostasis, PAI-1 is also involved, directly or indirectly through its action on plasminogen, in numerous biological processes including tissue remodelling, cell migration, and senescence [3,4]. PAI-1 is therefore widely regarded as a risk factor in various pathologies (cancer, cardiovascular diseases, age-related conditions), and several PAI-1 inhibitors are currently under development to suppress its activity [4]. While investigating the PAI-1 knockout model to study its effect in senescence [3], we observed Congo red-positive deposits in the kidneys of middle-aged animals. This finding prompted us to conduct an in-depth characterization of this model, with the dual goal of providing the scientific community with a novel animal model of amyloidosis and highlighting a potentially adverse effect of PAI-1 inhibition. All experimental procedures have been approved by our institutional review board for animal experimentation (APAFIS #33218-2021092417194431).

## Results

Congo-red positive amyloid deposits were observed in all PAI-1^−/−^ mice older than 12 months, affecting the spleen, kidney, liver, heart, tongue, and intestines (**Figure 1A, B and C**). No amyloid was detected prior to this age (**Figure 1C**). Wild-type controls from the same C57BL/6 genetic background (known to develop APOA2 amyloidosis in aged animals) showed significantly lower amyloid scores (p<0.001 in all organs) and penetrance (43% vs 100%) compared to PAI1^−/−^ mice, indicating an accelerated onset of amyloidosis in these animals (**Figure 1D**). To determine the type of amyloidosis, we performed mass spectrometry (MS) analysis on purified fibrils. Fibrils purification and characterization (**Supplementary Figure 1A and B**) and MS analysis were performed as previously described except that fibrils were digested with thermolysin instead of trypsin [5]. Western blotting of purified fibrils with a rabbit polyclonal anti-mouse APOA2 (ABClonal #A14690) revealed by an anti-rabbit IgG-HRP (cell signaling #7074P2) showed a major band at ~8 kDa (**Supplementary Figure 1B**) and MS identified peptides covering the full length secreted APOA2 protein as the main constituent of the deposits, including the propeptide known to be highly amyloidogenic (**Supplementary Figure 1C**) [6]. Low quantities of classical amyloid signature proteins such as APOE, COL6A2, COL6A1, VTNC, CLU, SAP and APOA4 were also detected (**Supplementary Figure 1D**). We confirmed APOA2 deposits by immunofluorescence on organs using the anti-mouse APOA2 antibody and a secondary antibody goat anti-rabbit FITC (Cell Signaling # 86426S) (**Figure 1E**). Sanger DNA sequencing of the APOA2 gene using primers 5’UTR-Fw_TGTTCCTAGGCCATAGTCTGC and 3’UTR-Rev_GAAGACTGGGCCTGGCAC, was consistent with the *APOA2*^*a*^ allele which is not associated with accelerated amyloidosis (**Supplementary Figure 1E**) [1] and mouse APOA2 dosage (ELISA Kit Biotechne #NBP2-68244) did not show any increase in serum APOA2 protein levels, which could have explained an earlier onset of the disease (**Figure 1F**). Similarly with previous systemic amyloidosis models [7], we showed that *i*.*v*. injection of Amyloid Enhancing Factor (AEF), prepared from the spleens of AAPOA2-positive PAI-1^−/−^ mice (0.1 mL), significantly accelerated the onset of amyloid deposits. 6 months old mice injected with one dose of AEF at 3 month-of-age were all positive, while only one was barely positive in the control group (spleen amyloid score: 1.93 ± 0.19 *vs* 0.08 ± 0.2, *p* = 0.0012) (**Figure 1G**).

**Figure 1.**
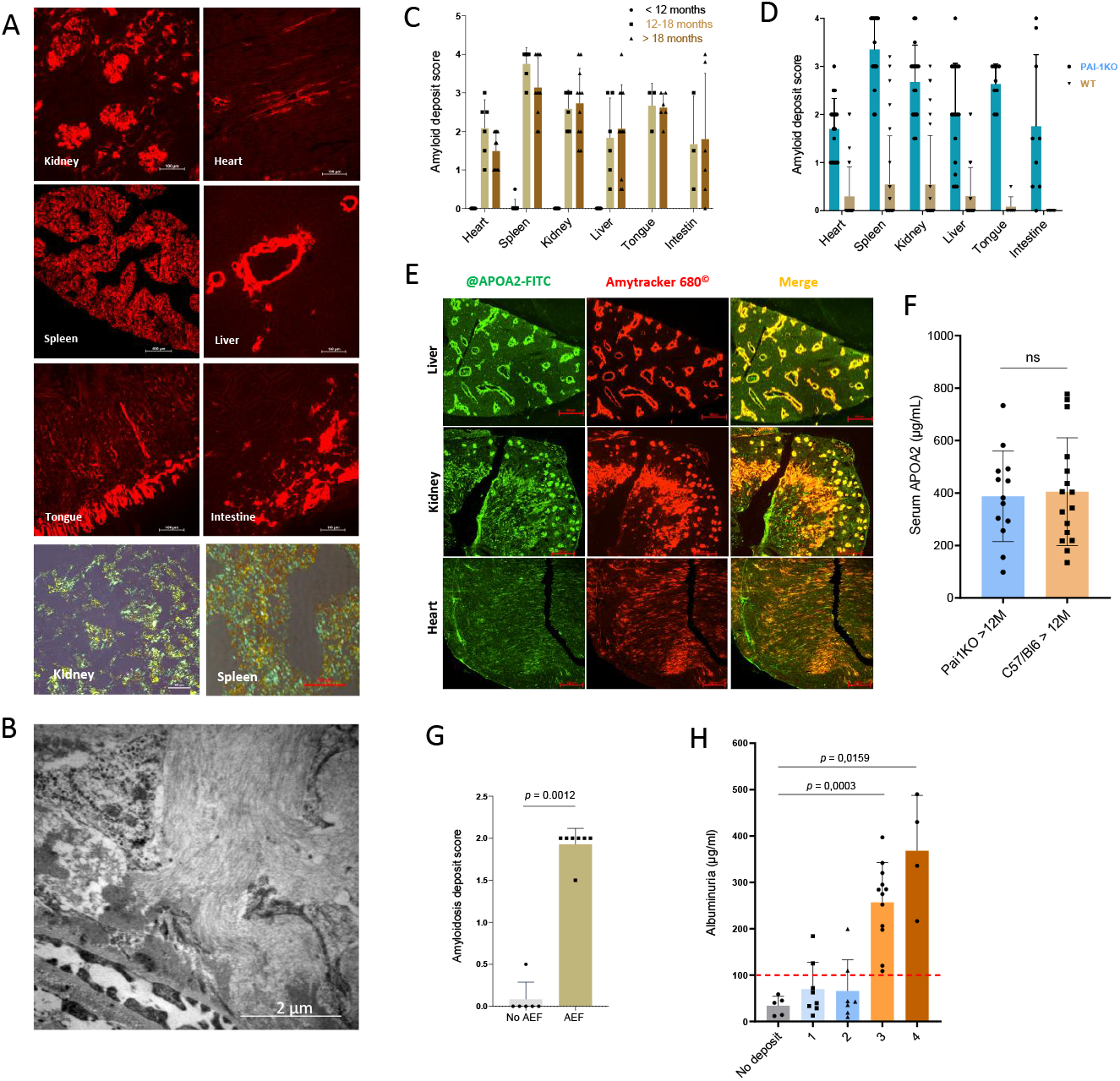
Characterization of senile APOA2 amyloidosis in PAI-1^−/−^ mice. (A) Congo-red staining in organs of PAI1^−/−^ mice. Representative images of Congo-red fluorescence in different organs are shown. Lower panel, Congo red birefringency in spleen and kidney of a PAI1^−/−^ mouse. Scale bars are indicated. (B) Electron micrograph of kidney from a PAI-1^−/−^ mouse showing amyloid fibrils in the extracellular compartment. Scale bars are indicated. (C) Amyloidosis scores relative to age in organs of PAI-1^−/−^ mice (n = 6-11 except tongue and intestine, n = 3-5). Protein deposition was graded on a four-point scale: 1, rare localized deposits; 2, frequent localized deposits; 3, deposits throughout the organ; and 4, deposits throughout the organ with visible tissue disorganization. (D) Amyloidosis scores in organs of > 12 months old PAI-1^−/−^ (n = 17) and C57Bl/6 control mice (n = 28). *p* < 0.0001 for all organs except intestine *p* = 0.013 (multiple t-tests, Benjamini–Hochberg FDR correction) (E) Representative immunofluorescence with an anti-mouse APOA2 antibody colocalized with Amytracker 680^©^ in organs of an 18 months old PAI-1^−/−^ mouse. Scale bars are indicated. (F) Serum APOA2 dosage in PAI-1^−/−^ (n = 13) and control mice (n = 16). (G) Amyloidosis scores in spleens of 6 months old PAI-1^−/−^ mice injected (n = 6) or not (n = 7) with AEF at month 3. (H) Urine albumin dosage relative to the amyloid score in spleens of PAI1^−/−^ mice (n ≥ 4 per score group). In F-H, differences between groups were calculated using a two-tailed Mann-Whitney test (Prism v8 GraphPad). *P* values are indicated when *p* < 0.05.

We next sought to determine whether APOA2 amyloid deposits induced organ dysfunction in PAI-1^−/−^ mice. Our goal was both to determine whether the intensity of amyloid deposition correlated with organ dysfunction and to identify a measurable marker that could predict amyloidosis in vivo. To this end, we measured two biomarkers commonly used to assess cardiac (serum NT-proBNP) and renal (albuminuria) dysfunction in amyloidosis, as previously described [5,8]. We found a significant correlation between albuminuria levels and the intensity of amyloid deposits (**Supplementary Figure 1F**). Particularly, in mice with high amyloid burden (score 3-4), an albuminuria level above 100□µg/mL reliably indicated the presence of amyloidosis (**Figure 1H**). We confirmed that albuminuria is linked to amyloid deposits rather than aging, as C57BL/6 WT mice older than 18 months showed no increase in urinary albumin, except for a single animal with a visibly abnormal kidney (**Supplementary Figure 1G**). In contrast, serum NT-proBNP levels did not correlate with cardiac deposits, consistent with the comparatively lower amyloid burden in the heart than in other organs (**Supplementary Figure 1H**).

## Discussion

We herein describe a novel murine model of accelerated apolipoprotein A-II (APOA2) amyloidosis, arising in plasminogen activator inhibitor-1 deficient (PAI-1^−/−^) mice. Unlike senescence-accelerated mouse strains or sporadic cases in aged mice, this model develops systemic amyloid deposits with complete penetrance at an intermediate age, thereby offering a robust and reproducible system for investigating age-related amyloidosis. The widespread distribution of deposits across multiple organs, together with the correlation between renal amyloid burden and albuminuria, underscores its potential utility for pre-clinical testing of therapeutic strategies aimed at preventing amyloid formation or removing deposits. The main limitation of our study is the absence of clear mechanistic explanation linking APOA2 amyloidosis to the absence of PAI-1. Few studies have reported such an association, generally suggesting that plasmin, increased by PAI-1 loss, inhibits amyloid development [9], in contrast to our findings. However, plasmin has been shown to play a direct role in ATTR amyloid formation by cleaving TTR and releasing a C-terminal fragment with high amyloidogenic potential [7]. This raises the possibility that increased plasmin activity due to PAI-1 inhibition could accelerate amyloidogenesis. Yet, in our model, full-length APOA2 was consistently identified as the fibril component. Thus, accelerated amyloidosis in PAI-1^−/−^ mice likely reflects indirect effects of PAI-1 loss. PAI-1 inactivation alters the extracellular matrix via plasmin-mediated activation of matrix metalloproteinases (MMPs), leading to enhanced degradation and remodeling [10]. Such changes may predispose tissues to the implantation of amyloid deposits. Further studies are needed to clarify whether this predisposition is specific to AApoA2 or applies to other systemic amyloidoses (ATTR, AL). Importantly, PAI-1, long considered deleterious in aging and fibrosis, is a target of pharmacological inhibitors under development [4]. Our findings suggest that chronic PAI-1 inhibition, while potentially beneficial, may inadvertently facilitate amyloid deposition and increase the systemic amyloidosis risk, an unanticipated concern in current clinical programs.

In conclusion, PAI-1^−/−^ mice provide a reproducible model of accelerated APOA2 amyloidosis with systemic involvement and accessible biomarkers for preclinical evaluation of anti-amyloid therapies or imaging tools. At the same time, they uncover an unexpected risk of PAI-1 inhibition, namely reduced tissue resistance to amyloid deposition. Whether this phenomenon extends to other forms of amyloidosis remains to be clarified, but it may offer novel insights into the interplay between extracellular matrix remodelling and protein instability in amyloid diseases.

## Abbreviations

PAI: plasminogen activator inhibitor
APOA-II: Apolipoprotein A-II
IF: immunofluorescence
AEF: Amyloid Enhancing Factor

## Acknowledgments

The authors thank the staff of the Biologie Intégrative Santé Chimie Environnement (BISCEm) technical platforms at the University of Limoges and Aurore Danigo from the pathology department of the University Hospital Dupuytren, Limoges, for her help on electron microscopy.

## References

[1] Korenaga T, Fu X, Xing Y, et al. Tissue distribution, biochemical properties, and transmission of mouse type A AApoAII amyloid fibrils. Am J Pathol. 2004;164(5):1597–1606.

[2] Carmeliet P, Kieckens L, Schoonjans L, et al. Plasminogen activator inhibitor-1 gene-deficient mice. I. Generation by homologous recombination and characterization. J Clin Invest. 1993;92(6):2746–2755.

[3] Cohen C, Le Goff O, Soysouvanh F, et al. Glomerular endothelial cell senescence drives age-related kidney disease through PAI-1. EMBO Mol Med. 2021;13(11):e14146.

[4] Khoddam A, Miyata T, Vaughan D. PAI-1 is a common driver of aging and diverse diseases. Biomed J. 2025;100892.

[5] Martinez-Rivas G, Ayala MV, Bender S, et al. A mouse model of cardiac immunoglobulin light chain amyloidosis reveals insights into tissue accumulation and toxicity of amyloid fibrils. Nat Commun. 2025;16(1):2992.

[6] Gursky O. Hot spots in apolipoprotein A-II misfolding and amyloidosis in mice and men. FEBS Lett. 2014;588(6):845–850.

[7] Slamova I, Adib R, Ellmerich S, et al. Plasmin activity promotes amyloid deposition in a transgenic model of human transthyretin amyloidosis. Nat Commun. 2021;12(1):7112.

[8] Bender S, Ayala MV, Bonaud A, et al. Immunoglobulin light-chain toxicity in a mouse model of monoclonal immunoglobulin light-chain deposition disease. Blood. 2020;136(14):1645–1656.

[9] Liu R-M, van Groen T, Katre A, et al. Knockout of plasminogen activator inhibitor 1 gene reduces amyloid beta peptide burden in a mouse model of Alzheimer’s disease. Neurobiol Aging. 2011;32(6):1079–1089.

[10] Flevaris P, Vaughan D. The Role of Plasminogen Activator Inhibitor Type-1 in Fibrosis. Semin Thromb Hemost. 2017;43(2):169–177.

