## Supplementary material for "Accelerated APOA-II senile amyloidosis in a *PAI-1* (plasminogen activator inhibitor 1) knock-out model": supp Figure 1

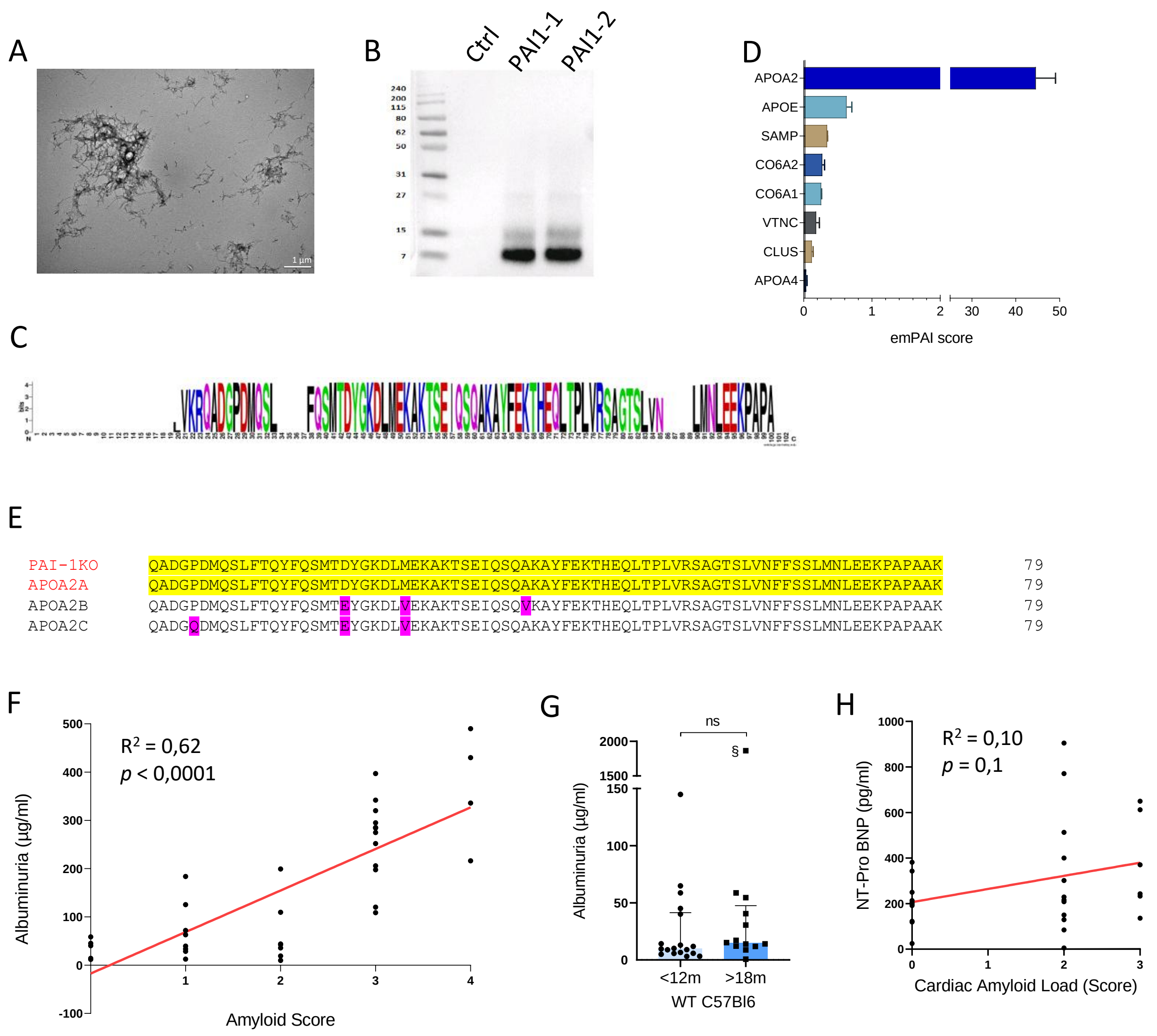


**Supplementary Figure 1.** (A) Electron micrograph of purified fibrils from a PAI-1$^{-/-}$ mouse spleen. Scale bars are indicated. (B) Analysis of the purified fibrils by Werstern blot using a anti-APOA2 antibody. Ctrl, spleen from a C57BL/6 mouse without amyloid deposits (C) Weblogo representation of the APOA2 aminoacid sequence coverage upon MS analysis. Purified fibrils were digested by thermolysine and 2 µL of the resulting digestion was injected in a Bruker TimsTof 2 device. (D) Relative abundance (EmPAI score) of APOA2 and other amyloid signature proteins in purified fibrils extracted from PAI-1$^{-/-}$ mice (n = 3) (E) Comparison of APOA2 deduced protein sequence found in PAI-1$^{-/-}$ with known APOA2 isoforms showing a full homology with APOA2$^{a}$ (highlighted in yellow). The position of amino acids defining the different isoorms are highlighted in purple. (F) Simple linear regression showing the correlation between renal amyloid burden and albuminuria. $R^2$ and $p$ value are indicated. G. Albuminuria is not linked to age in C57BL/6. n = 18 in < 12 months mice and n = 13 in > 18 months mice. Two-tailed Mann-Whitney test (Prism v8 GraphPad). § This mouse presented with abnormal pale kidneys at sacrifice. (H) Simple linear regression showing the absence of correlation between cardiac amyloid burden and serum NT-proBNP levels. $R^2$ and $p$ value are indicated.
